# Modeling Protein Conformations by Guiding AlphaFold2 with Distance Distributions. Application to Double Electron Electron Resonance (DEER) Spectroscopy

**DOI:** 10.1101/2024.10.30.621127

**Authors:** Tianqi Wu, Richard A. Stein, Te-Yu Kao, Benjamin Brown, Hassane S. Mchaourab

## Abstract

We describe a modified version of AlphaFold2 that incorporates experiential distance distributions into the network architecture for protein structure prediction. Harnessing the OpenFold platform, we fine-tuned AlphaFold2 on a small number of structurally dissimilar proteins to explicitly model distance distributions between spin labels determined from Double Electron-Electron Resonance (DEER) spectroscopy. We demonstrate the performance of the modified AlphaFold2, referred to as DEERFold, in switching the predicted conformations guided by experimental or simulated distance distributions. Remarkably, the intrinsic performance of AlphaFold2 substantially reduces the number and the accuracy of the widths of the distributions needed to drive conformational selection thereby increasing the experimental throughput. The blueprint of DEERFold can be generalized to other experimental methods where distance constraints can be represented by distributions.

## Introduction

AlphaFold2^1^ has revolutionized the field of protein structure prediction. Long considered the holy grail of the protein folding problem^2–5^, the decoding of the sequence-structure relationship emerged from the training of a deep learning neural network model^1,6–10^ on the wealth of protein sequences^11^ and high-resolution structures^12^. The accuracy of AlphaFold2 models revolutionized structural biology by providing interpretive frameworks for classic structure/function studies of proteins, starting models for cryo-electron microscopy (cryo-EM) and X-ray crystallography^13–15^, and powerful tools for in silico drug screening^16,17^.

Its performance notwithstanding, the success of AlphaFold2 belied two fundamental limitations that are front and center in the field. The first limitation arises from the fact that the quality of its models depends on the depth and information content of the multiple sequence alignment (MSA)^1,18^. Hence, model prediction from a single sequence is challenging, which drove the development of alternative deep learning methods^19^. The second limitation arises from the assumption that an input sequence is associated with a single model. The latter ignores the rugged conformational landscape of protein folded states deduced from experimental data and supported by Molecular Dynamics (MD) simulations^20,21^. The meandering of proteins between conformations, driven by thermal energy, or by triggered large-scale conformational changes underpins biological function. Multiple methods have been advanced to “coerce” AlphaFold2 to generate multiple conformations or protein ensembles through modifications of the MSA^22–24^. While these methods of “hacking” AlphaFold2 have achieved remarkable success, their performance appears to be target-dependent.

Experimental structural biology techniques have been developed to probe protein conformational ensembles providing snapshots along the energy landscape. While the recent emergence of single particle cryo-EM facilitated direct visualization of intermediates in some cases, insight into protein conformational dynamics has been the forte of a battery of biophysical techniques such as nuclear magnetic resonance (NMR)^25^, probe-based spin labeling EPR spectroscopy^26–29^ and fluorescence resonance energy transfer^30,31^, as well as hydrogen-deuterium exchange^32^ and cross-link mass spectrometry^33,34^. Observables from these techniques are formulated as constraints that are dependent on the biochemical conditions that modulate conformational sampling.

Since the early days of AlphaFold2, there has been an emphasis on the experimental validation and subsequent refinement of its predictions^15,22,23,35–37^. With the availability of AF2 methods to generate conformational ensembles, the goal is to place the models in the context of the energy landscape. Methods to incorporate the experimental data from biophysical and biochemical measurements into the prediction engine of AlphaFold2 would streamline such a process and provide a general tool for iterative refinement. AlphaLink^37^ was an early pioneering example that fine-tuned AlphaFold2 with data obtained from cross-linking mass spectrometry. Despite its limitations to one modality, it showcased the flexibility of the network to experimental input although the question of constrained predictions of multiple conformations was not addressed. We explore this aspect of AlphaLink here.

In addition, and more generally, the incorporation of constraints, particularly from probe-based spectroscopic techniques, presents a unique challenge due to the rotameric freedom of the probe relative to the backbone. For example, Double Electron Electron Resonance (DEER) spectroscopy^26,38^ reports a distance histogram between two spin label sites. There is a rich history of modeling the transfer function between the spin label distances and the corresponding alpha carbon distances^39–44^. Yet, all the models share an inherent uncertainty in modeling the spin label rotameric freedom. This uncertainty has hampered the interpretation of the absolute distance and can mask the distance changes that result from conformational rearrangements, one of the strengths of DEER spectroscopy. Similar issues confront the applications of Fluorescence Energy transfer.

In this work, we develop an AlphaFold2 method that overcome the challenges of incorporation of DEER distributions and provide a blueprint for extension to other experimental spatial constraints expressed as distributions. Testing AlphaLink shows that it can drive modeling alternative conformations from distance distributions between alpha carbons. AlphaLink’s predictions, however, show sensitivity to the width of distance distribution reinforcing the need for explicit consideration of the spin label side chains into the network architecture. Therefore, we finetuned AlphaFold2, in the context of OpenFold^45^(a trainable PyTorch reproduction of AlphaFold2), to interpret spin-label distance distributions determined from DEER spectroscopy and integrate them into the network architecture resulting in a new method named DEERFold (**Fig. 1**). We conducted a systematic evaluation of DEERFold’s performance on a diverse set of water soluble and membrane proteins. Our results demonstrate that including experimental or simulated spin label distance constraints drives model folding into the desired conformation. Optimization of the information content of the constraints improve the quality of the predictions. Remarkably, we found that the exact shape of the distributions did not influence the ability of DEERFold to model the targeted conformation. For general applicability, we develop an approach to address evaluate DEERFold results in the absence of initial and target experimental structures.

**Fig. 1:**
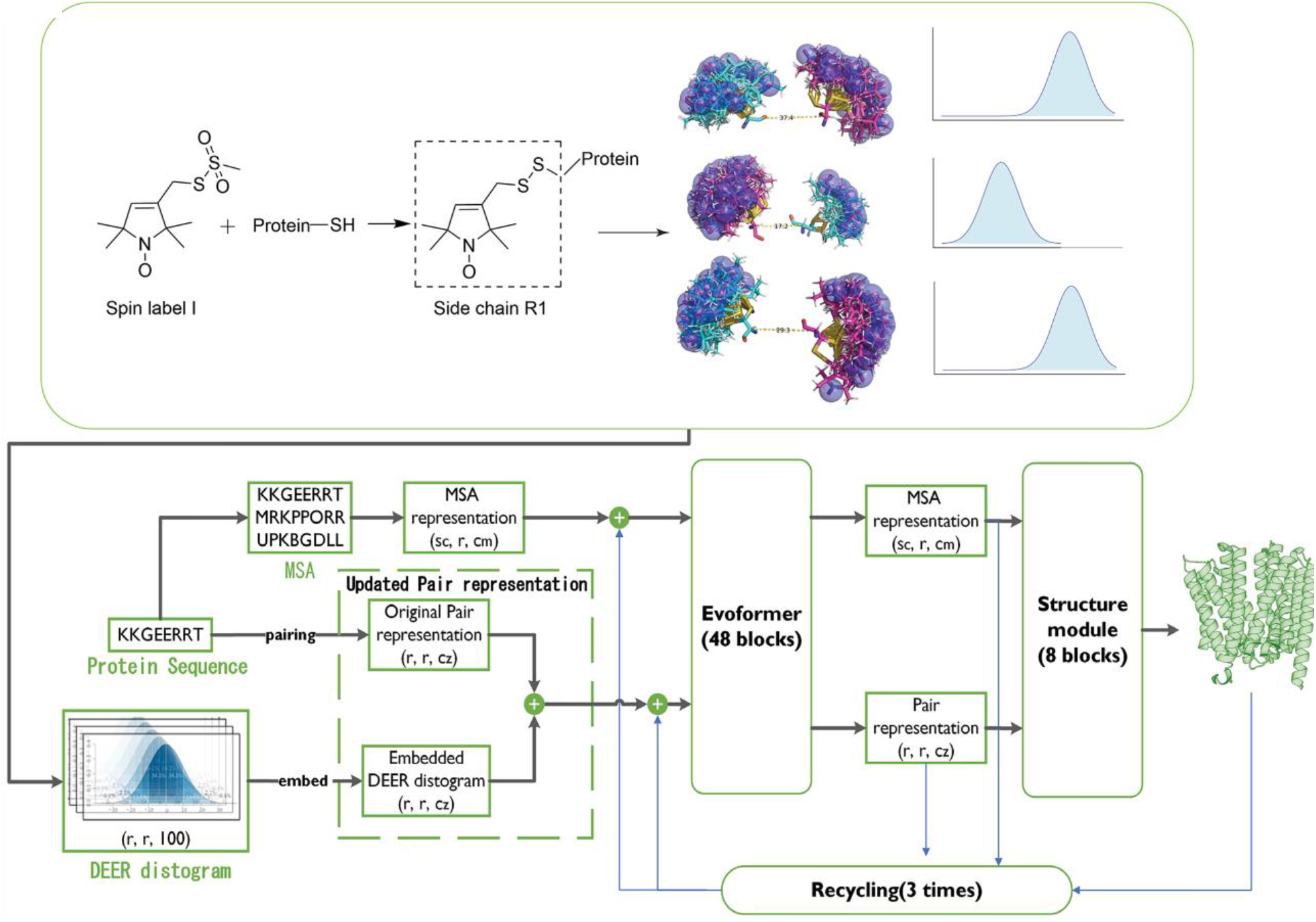
DEERFold architecture showing the DEER distogram input.

## Methods

### Overview of DEERFold

To fully leverage the potential of DEER data in protein structure prediction, we have developed DEERFold, a framework that directly incorporates spin label distance distributions derived from DEER data into AlphaFold2 through OpenFold^45^. OpenFold is a trainable PyTorch reproduction of AlphaFold2, and its developers have verified that the implementation yields result comparable to those of the original AlphaFold2^45^.

AlphaFold2 primarily exploits co-evolutionary relationships. The main challenge in merging multiple information sources is to find a suitable representation that facilitates integration while avoiding information loss. AlphaFold2 operates both in coevolution space (Evoformer) and 3D space (Structure Module). Spin label distances derived from DEER data align well with AlphaFold2’s distance space. Thus, we integrated the DEER data into AlphaFold2 via the Evoformer, complementing existing co-evolutionary data within AlphaFold2 to improve the prediction accuracy and predict different protein conformations.

### Model Architecture and Training Strategies

The total dataset comprised 6,565 protein chains with sequence identity ≤ 30% using MMseqs2^46^. Of these, 6,202 were used for training randomly sampled from the OpenFold dataset (https://registry.opendata.aws/openfold/)^45^. For validation, we used a manually curated dataset^47^ of 363 proteins, consisting of open-closed conformational pairs and their intermediate states as defined by the original authors of each PDB entry. For each chain, potential spin labeling sites were first identified as helical or beta strand, and solvent-exposed residues by DSSP^48^. Pairwise spin label distance distributions for labeling site pairs larger than 15Å were calculated using chiLife^42^.

The input features to AlphaFold2 consisted of the spin label distograms as pairwise residue restraints, along with the pre-computed multiple sequence alignments (MSAs) provided in OpenFold. The distogram shape was LxLx100, with each bin, except for the first and last, spans 1 Å and is centered at integer values from 1 to 99 Å. The first bin includes all values up to and including 1.5 Å, while the last bin captures all values above 99.5 Å. Targets in OpenFold with sequences mismatched to the MSA were filtered out. For each epoch, we randomly selected up to 25 pairs of spin label sites per target. Following the AlphaLink training strategy, a random MSA subset with Neff^49^ between 1-25 was sampled per epoch and target.

For AlphaLink testing, input distance constraints are formatted as Cα-Cα distances, represented by a 128-bin distogram ranging from 2.3125 Å to 42 Å. The input distance constraints were converted to match AlphaLink’s original format.

The overall model architecture (**Fig. 1**) integrates the AlphaFold2 framework with spin label distograms as an additional pairwise input feature channel in the Evoformer module. The Evoformer integrates the multiple sequence alignment (MSA) representation (dimensions: (s_c_, r, c_m_), where s_c_ is the number of randomly selected sequences, r is the query sequence length, and c_m_ = 128) and the pair representation ***z*** (dimensions: (r, r, c_z_), where c_z_ = 128), providing structural information to the Structure Module. The MSA representation encodes individual residues and their homologous sequences, while the pair representation captures pairwise relationships between residues. DEER data, in the form of distance distributions (100 bins), is incorporated into the pair representation by replacing template information. The Evoformer block processes the updated pair representation ***z***, which includes DEER distance information, allowing the MSA and pair representations to exchange information biased towards the DEER distogram data.

Implementation was based on the retrainable PyTorch version of AlphaFold2 from the OpenFold package^45^. The model was fine-tuned using AlphaFold2’s pre-trained model weights (model_5_ptm) that don’t use templates and follows AlphaFold2’s fine-tuning process. We used the previously released version of AlphaFold2 trained using PDB structures with a release date before 2018-04-30. We employed the Adam optimizer^50^ with an initial learning rate of 0.0005. The fine-tuning was performed in a data-parallel fashion across 4 NVIDIA A6000 GPUs (48 GB memory each), with one protein chain distributed per GPU. A held-out validation set of 962 chains from the OpenFold dataset was utilized for model selection. We systematically benchmarked the trained model’s performance on a diverse set of proteins, including AK(open conformation:4AKE^51^; closed conformation:1AKE^52^), RBP(open conformation:1BA2^53^; closed conformation: 2DRI^54^), and a set of membrane protein targets including the proton-coupled multidrug transporter PfMATE(outward-facing conformation:6GWH^55^; inward-facing conformation:6FHZ^55^), the major facilitator superfamily antiporter LmrP(outward-facing conformation:6T1Z^56^; inward-facing conformation: AlphaFold2_prediction^57^), the multidomain ABC transporter ABCB1 or P-glycoprotein (Pgp) (outward-facing conformation:7ZK4^58^; inward-facing conformation:7A65^59^,7ZK7^58^), MCT1(inward-open conformation:7DA5^60^; outward-open conformation:7CKR^60^), STP10(outward occluded conformation:7AAQ^61^; inward open conformation:7AAR^61^), LAT1(inward open conformation:6IRS^62^; outward-facing occluded conformation:7DSQ^63^), ASCT2(inward-open conformation:6RVX^64^; outward-open conformation:7BCQ^65^), E. coli SemiSWEET(inward conformation: 4×5M^66^; outward conformation: 4×5N^66^), and DgoT (inward conformation:6E9N^67^; outward conformation:6E9O^67^), which had been previously characterized to adopt distinct conformational states. To assess whether the model biases towards particular structural folds, an additional model was trained on the same training data but using spin label pairs on the surface sites of β-strands.

Neff^49^ was calculated by assigning a weight to each sequence in the multiple sequence alignment (MSA). A sequence’s weight was determined by how many other sequences in the MSA were similar to it, using a similarity cutoff of 0.62. Sequences with many similar counterparts received lower weights. Neff was then computed as the sum of these weights, effectively reducing the impact of redundant sequences in the alignment. In this work, the MSAs for those targets were set to Neff = 5 and generated by MMesqs2^68^ searching against ColabFold databases^18^(uniref30_2302).

#### Root Mean Square Deviation (RMSD)

RMSD is a commonly used metric to quantify the structural similarity between a predicted model and a reference model. In this study, we calculate the RMSD between the corresponding Cα atoms of the predicted and reference structures. The RMSD is defined as:

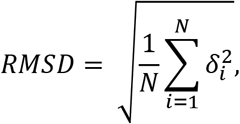

where 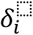 is the distance between Cα atom *i* in a predicted structure and a reference structure, and *N* is the number of Cα atoms being compared at the positions of interest. A lower RMSD value indicates a higher degree of structural similarity between the predicted and reference models, with an RMSD of 0 Å corresponding to identical structures at the compared positions.

### Root Mean Square Error (RMSE)

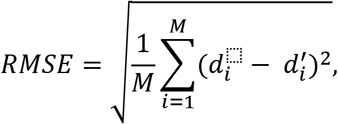

where *M* is the number of residue pairs used as DEER distance constraints, 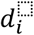 is the distance between the *i*-th residue pair in the predicted model, and 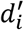 is the distance between the *i*-th residue pair in the reference model. In this study, we employed RMSE for the preliminary analysis conducted on Alphalink. For fair evaluation, we use the spin label pairs reported in published papers^27,28^, where high-resolution structural information for one or both conformational states was available to assess the model’s performance.

#### Template Modeling Score (TM-score)

TM-score^69,70^ is a metric used to assess the structural similarity between two protein structures, typically a predicted model and a native (reference) structure.

The TM-score is defined as:

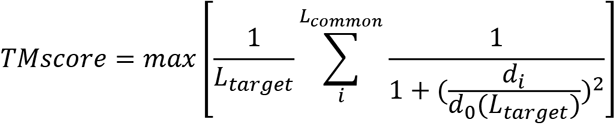

where *L*_*target*_ is the length of the target protein, and *L*_*common*_ is the number of the common aligned residues in two structures. *d*_*i*_ is the distance between the ith pair of residues in the template and target structures, and 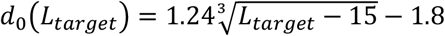 is a normalization factor that accounts for the protein’s length.

The TM-score ranges from 0 to 1, with a value of 1 indicating a perfect match between the two structures. A TM-score > 0.5 generally indicates that the two structures have the same fold, while a TM-score < 0.17 suggests that the similarity is likely due to random chance. It is less sensitive to local structural errors and more sensitive to global structural similarity, compared to RMSD.

### Earth Mover’s Distance (EMD)

Also known as Wasserstein distance^71^, EMD is a metric used to quantify the dissimilarity between two probability distributions. In the context of comparing spin-label distance distributions, EMD measures the minimum amount of “work” required to transform one distribution into the other, where “work” is defined as the product of the amount of probability mass moved and the distance it is moved. For each pair of spin labels, we first calculate the distance distribution from the DEERFold prediction using chiLife^42^, converting it to a discrete distribution over 100 distance bins ranging from 1 to 100 Å. To focus on the primary distance peak, we extract a 16-bin window centered on the maximum probability (±8 bins). The EMD is computed between the cumulative distribution functions (CDFs) of this processed prediction distribution and the input constraints. The final EMD score for a model is calculated as the mean EMD across all spin label pairs.

### Experimental and Simulated DEER Data

We evaluated DEERFold using both experimental and simulated DEER distance data. Experimental data, sourced from previous studies on membrane proteins solubilized in detergent micelles or lipid nanodiscs, were represented in two ways: **Experiment 1**(Multi-Gaussian fitting), where distance distributions were analyzed by fitting 2-4 Gaussians to the raw data, and **Experiment 2**(Single-peak approximation), where the highest-weight component corresponding to the from the multi-Gaussian fit was selected to create a single-peak distribution (std = 2 Å) to represent a more defined distance distribution. For LmrP and PfMATE, distance distributions at pH 8/7.5 and pH 4 were used for OF and IF predictions, respectively. For Pgp, distributions in the absence and presence of ADP, vanadate, and verapamil were used for IF and OF predictions, respectively.

Simulated distributions were generated using the optimization method of Kazmier et al.^72^, which maximizes information content for *ab initio* Rosetta folding. The method selects pairs of residues at the ends of long helix segments (≥ 20/30 residues) or beta strand segments (≥ 5 residues). Two sets of simulated distributions were created, and the corresponding distance distributions simulated using the MMM module in the chiLife^42^ package. **Simulation 1** employed a helix segment length threshold of 20 residues, while **Simulation 2** used a threshold of 30 residues. Each set yielded two spin-label distance distributions per helix/helix, or strand/strand segment pair.

## Results

We initially selected two transporters to test the hypothesis that spin label distances derived from DEER data could serve as direct input to guide AlphaFold2’s prediction process. The multidrug transporter LmrP^56^ is a proton-coupled antiporter of the major facilitator superfamily. It was one of the targets in CASP14^73^ and showcased the ability of AlphaFold2 to predict inward-facing (IF) and outward-facing (OF) conformations^73^. LmrP’s proton-powered conformational cycle was investigated by DEER in detergents and nanodiscs^28,29^ providing a set of real-world distance distributions for benchmarking DEERFold. Similarly, the second transporter, PfMATE^55^, is a proton-coupled multidrug transporter but from the multidrug and toxic compound extrusion (MATE) superfamily. Its crystal structures were determined in the IF and OF conformations^55^. An extensive DEER data set^27^ was collected for this transporter that monitored the proton-dependent conformational switching.

Our investigation proceeded in two phases. Initially, we took advantage of AlphaLink^33^ to test the conceptual feasibility of generating restrained AlphaFold2 models. AlphaLink is a version of AlphaFold2 fine-tuned on photo amino acids (photo-AA) crosslink constraints (simply referred to Cα-Cα contacts between photo-amino acids that are not within the same or consecutive tryptic peptides) as input to model 3D structures. Here we show its applications and limitations for constraints consisting of distance distributions between alpha carbons. These tests guided the refinement of AlphaFold2 to incorporate DEER distance distributions in the generation of models.

### Can AlphaLink be biased by distance distributions to generate alternate conformations

To test AlphaLink as a potential platform for incorporation of DEER constraints, we modeled input distance distributions as Gaussian with a standard deviation of 2 Å to reflect either rotameric distributions of the side chain or conformational flexibility of the backbone. We were particularly interested in testing if such a distance format can guide AlphaLink away from the default structure. For this purpose, we analyzed performance with three different sets of inputs. **Set 1**: Cα-Cα distances for all residue pairs, {*d*_*ij*_(*target*)| (*i, j*) ∈ [1, *L*], *L*: protein sequence length, *target*: target conformation}. **Set 2**: a reduced set of Cα-Cα distances involving strategically selected distance constraints that captures differences between the initial and target conformations {*d*_*ij*_(*target*)| (*i, j*) ∈ |*d*_*ij*_(*target*) – *d*_*ij*_(*initial*)| ≥ 10 Å, *nintna* : AlphaFold2 initial conformation, *target*: target conformation}. **Set 3**: Further refined constraints by focusing on regions with high AlphaFold2 pLDDT confidence scores {*d*_*ij*_(*target*)| (*i, j*) ∈ |*d*_*ij*_(*target*) – *d*_*ij*_(*initial*)| ≥ 10 Å, *plddt*_*i*_ ≥ 90, *plddt*_*j*_ ≥ 90}.

For PfMATE, the initial unconstrained AlphaFold2 prediction favored the OF structure, as shown in **Fig. 2A** (model in light blue on the left). Using the full set of Cα-Cα distances between all residue pairs in the IF structure as input (**Set 1**) we obtained a model that deviated substantially from both the OF and target IF conformations **(Fig. 2)**. The RMSE value was 16.90 Å compared to the initial OF model and 14.35 Å to the target IF model, with significant unresolved loop regions in certain areas.

**Fig. 2:**
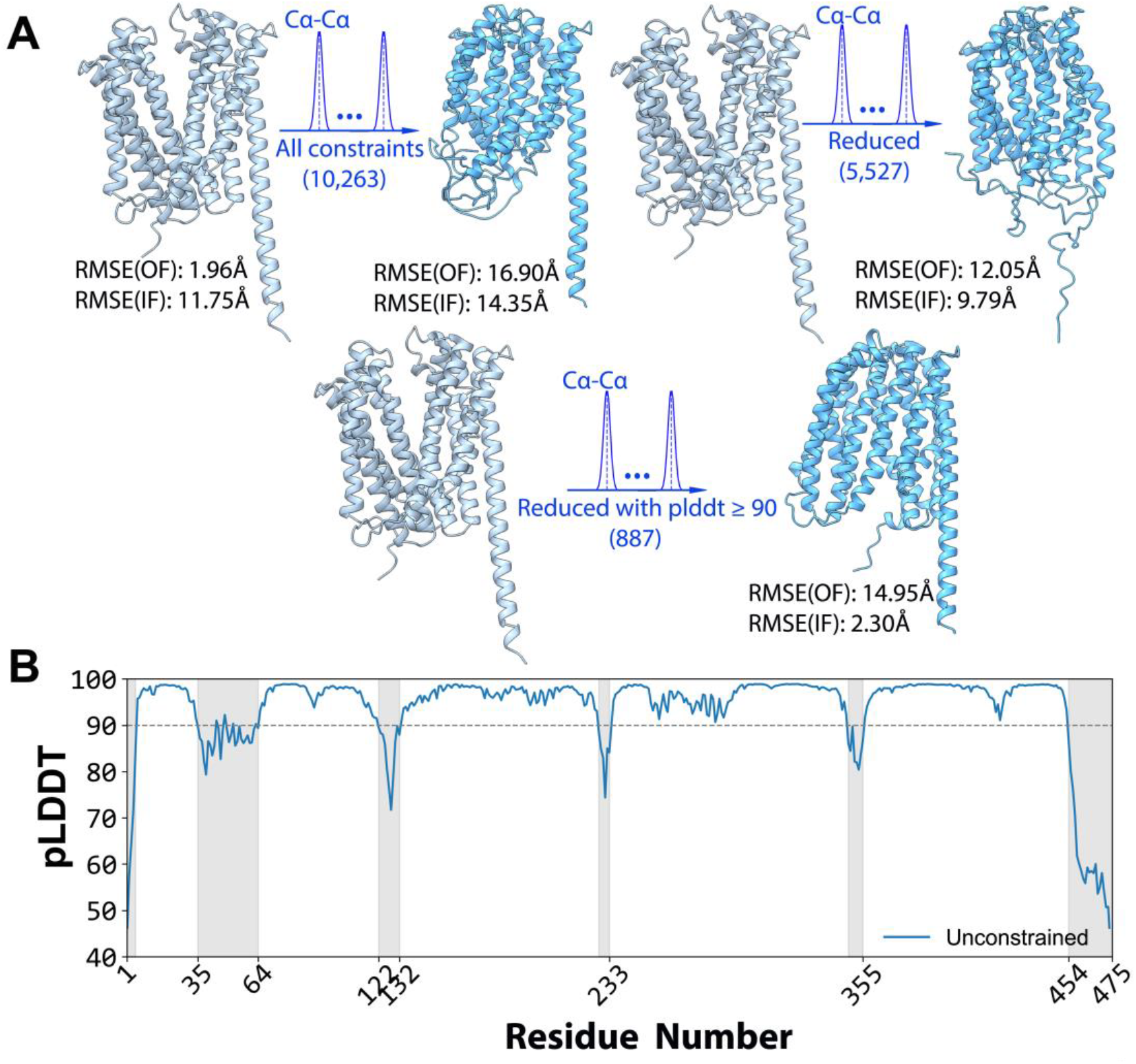
Impact of distance constraints on AlphaLink prediction of the target conformation for PfMATE. **A**. Initial unconstrained AlphaFold2 prediction favors the OF conformation in light blue and the predicted models from different constraint sets are shown in dark blue. The models were predicted using **Set 1**, all Cα-Cα distances (top left), **Set 2**, the subset of 5,527 distances from regions with structural differences (top right) and **Set 3**, further refined distance constraints from high pLDDT score regions (bottom). **B**. Per-residue pLDDT (predicted Local Distance Difference Test) score plot from the PfMATE unconstrained model, with the gray region containing low-confidence areas (pLDDT < 90).

We reasoned that the distorted model may be the result of an excessive number of distance constraints that overly restricted the model’s conformational sampling ability or included mutually incompatible constraints. Therefore, we reduced the input distances to a subset of 5,527 constraints from regions exhibiting significant structural differences between the target IF and initial OF states (**Set 2**). However, even with this reduction from the full 10,263 distances, the predicted model in **Fig. 2A** still deviated significantly from both conformations, with unstructured backbones in certain regions.

Building on this observation, we tested whether the failure of these models in recapitulating the target conformation may arise from a particular subset of constraints. Therefore, we compared unstructured regions in the unconstrained model against the target structure. We found that the pLDDT confidence scores for some regions from the initial unconstrained predictions are below 90 in the ranges [1,5], [35,64], [122,132], [228,233], [348,355], and [454,475] (the gray region in **Fig. 2B**). We reasoned that inaccurate distance restraints in these low-confidence areas were misguiding the folding process.

Therefore, we further refined the input by filtering out distance constraints from regions with pLDDT scores < 90, reducing the final constraint set from 5,527 to 887 distances (**Set 3**). With this refined, high-confidence subset of distances, AlphaLink demonstrated its ability to successfully fold towards the target IF conformation based solely on the Cα-Cα restraints derived from the known IF structure (**Fig. 2A**).

Similar findings were observed for LmrP as shown in Supplementary **Fig.1**. The analysis for PfMATE and LmrP demonstrates that a curated subset of input distance distributions, derived from regions predicted with high accuracy effectively guided AlphaFold2 towards the conformational states represented by those distances, in contrast to the unconstrained predictions or when using full distance sets that included low-confidence regions.

### The Impact of Distance Distribution Width on AlphaLink prediction

To test the effects of the width of the distance distributions, we selected a subset of input distance constraints from set 3 above that met two criteria: **1)** regions exhibiting significant structural differences between initial and target states within certain cutoffs, and **2)** pLDDT scores ≥ 90. While in **Fig. 2** and Supplementary **Fig.1, a** cutoff of 10 Å (|*d*_*ij*_(*target*) – *d*_*ij*_(*initial*)| ≥ 10 Å) was used, here we systematically evaluated different cutoff values. For each prediction (**Fig. 3** for PfMATE), we determined the RMSEs between predicted models using different distribution widths (std values) and reference structures – i.e. the default OF state(6GWH^55^) or the target IF(6FHZ^55^). We then calculated the average RMSEs (curve in **Fig. 3**) to the initial and target conformations across the 5 predicted models per std value.

**Fig. 3:**
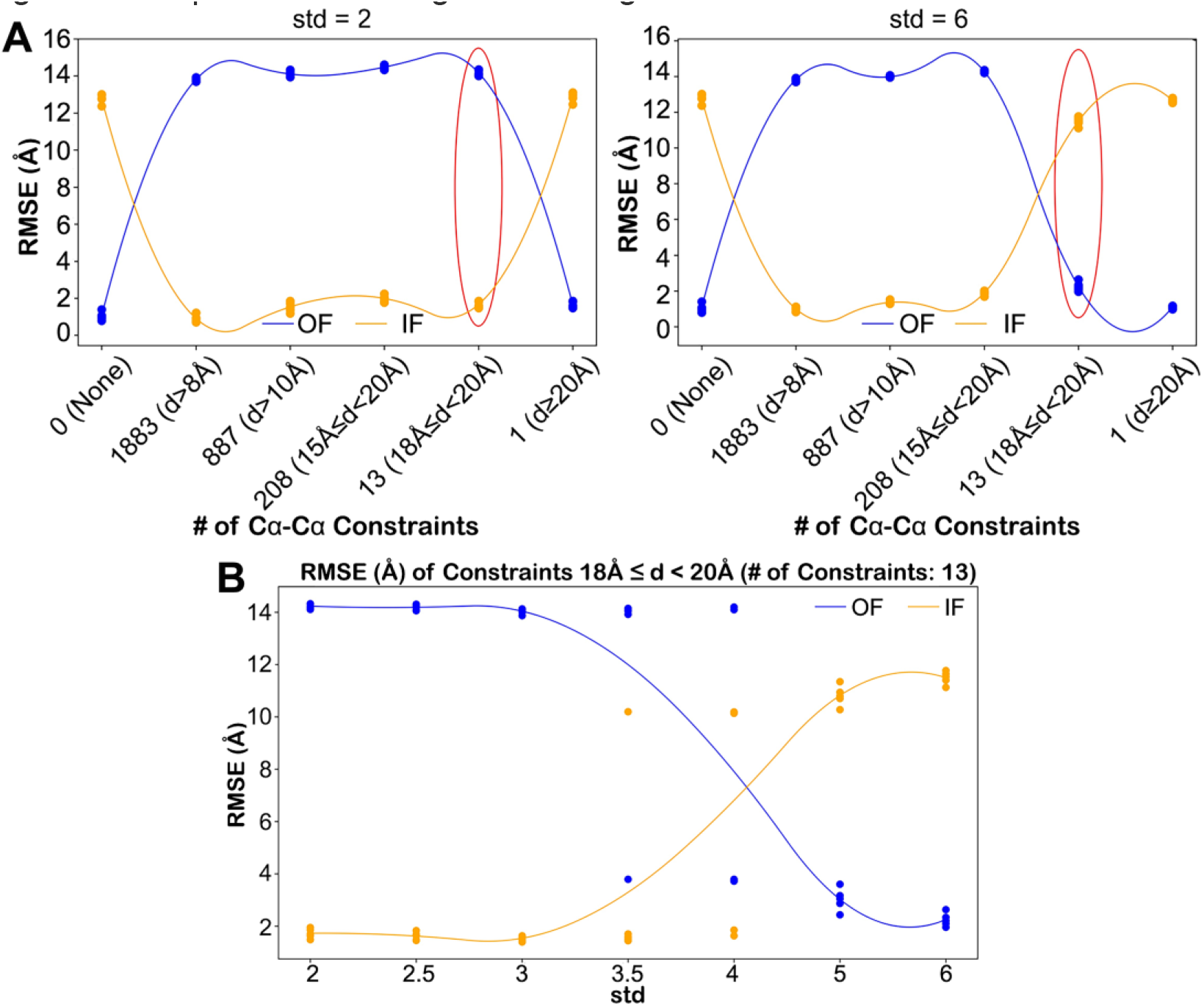
Impact of distance distribution widths on AlphaLink prediction of the target conformation for PfMATE. **A**. Effect of the number of distance constraints on prediction at std=2Å and std = 6 Å. **B**. With the same set of data at the cutoff [18 Å, 20 Å), AlphaLink prediction results from std=2 to std = 6 Å (The blue data points represent RMSEs of the predicted models relative to the OF state, while the yellow points indicate the RMSEs to the target IF state). The two curves show average RMSEs to initial(blue) and target(yellow) states across the ensemble of 5 predicted models per std value.

Remarkably in **Fig. 3A**, at std = 2Å, the limited set of constraints from the IF conformation (highlighted by the red circle) using the cutoff {*d*_*ij*_(*target*)| (*i, j*) *ϵ* 18Å ≤ |*d*_*ij*_(*target*) – *d*_*ij*_(*initial*)| < 20Å)] still guided AlphaLink towards the target IF conformation, away from the initial unconstrained OF model. In contrast, at the width of std = 6 Å, the same set of distance distributions failed to recover the IF state, suggesting that a tipping point lies between 2-6 Å for this cutoff set.

In **Fig. 3B** with the same set of data, std values of 2, 2.5, and 3 Å, all effectively guided AlphaLink into the expected IF conformation. But at std = 3.5 Å, the 5 models displayed considerable variability, with one model folding into the OF conformation and four models predicting the IF conformation. At std = 4 Å, only 2/5 models satisfied the constraints. Finally, extending to 6 Å std prevented folding into the target IF conformation based on these distances.

Similar to PfMATE, we analyzed full-distance cutoff sets at std = 2 Å and 6 Å to identify a critical cutoff range for LmrP. From **Fig. 4A**, we observed that even the rather strict [15 Å, 20 Å) cutoff, containing only 29 Cα-Cα residue pairs, could still guide AlphaLink folding to the target state at 6 Å std. In contrast to PfMATE, std values from 2 Å up to 6 Å all successfully folded the unconstrained starting model into the final IF state shown in **Fig. 4A**. Narrowing the distributions below 2 Å std in the cutoff {*d*_*ij*_(*target*)| (*i, j*) *ϵ* |*d*_*ij*_(*target*) – *d*_*ij*_(*initial*)| ≥ 8Å)], which yielded 2,063 total constraints led to an unexpected observation. In the extreme limit of std = 0 Å, representing a single fixed distance, the predicted models appeared trapped in an intermediate state between the initial and target, with average RMSEs of 4.36 Å and 3.74 Å respectively (**Fig. 4B**). However, even a slight increase to 0.5 Å std enabled the models to successfully satisfy the input distances and fold into the target conformation. This highlights how overly narrow distance distributions albeit with a large input set can restrict AlphaLink from sampling the correct conformation.

**Fig. 4:**
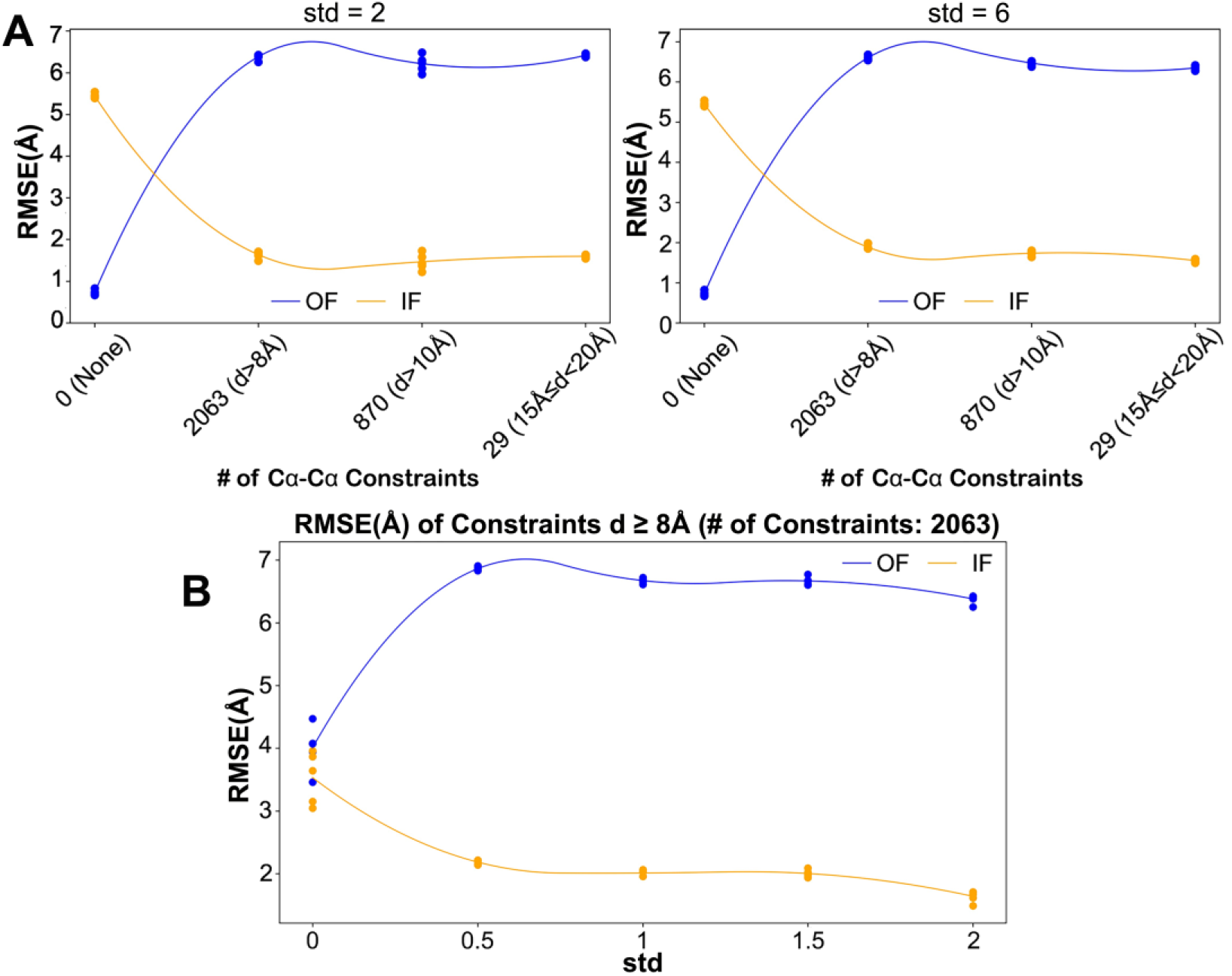
Impact of distance distribution widths on AlphaLink prediction of the target conformation for LmrP. **A**. Effect of distance cutoffs on prediction at std=2Å and std = 6 Å. **B**. With the same set of data at the cutoff [8 Å, +∞), AlphaLink prediction results from std=2 to std = 6 Å (The blue data points represent RMSEs of the predicted models relative to the OF state(6T1Z), while the yellow points indicate the RMSEs to the target IF state). The two curves show average RMSEs to initial(blue) and target(yellow) states across the ensemble of 5 predicted models per std value.

Overall, our feasibility testing establishes that distance constraints between alpha carbons modeled as distributions, rather than fixed values, can guide AlphaFold2 to sample multiple conformations. In fact, distance constraints, formulated as a single fixed distance, may lead to prediction failures by excessively limiting the sampling of conformational space. Conversely, the range of standard deviations from 2 to 3 Å appears to allow sufficient variability while still effectively guiding the folding process for both PfMATE and LmrP. Broader distributions fail in general suggesting that for such distributions, AlphaFold2 needs to be refined/trained with appropriate widths.

### Spin Label Distance Distributions fail as input for AlphaLink

Initially, we attempted to directly input the experimental spin label distances into AlphaLink by converting them into distribution representations (distograms with shape LxLx128, comprising 127 distance bins spanning 2.3125-42Å at 0.3125Å intervals, and a catch-all bin for >=42Å). However, this approach yielded unsatisfactory predictions, revealing a wide range of errors between the spin label distances and the actual Cα-Cα distances within the protein structures.

For PfMATE (**Fig. 5A**), we selected residue pairs to generate input distance restraints from the experimental spin label data reported previously^27^. Following the guidelines from the tests above, we modeled the spin label distances as LxLx128 distributions using chiLife^42^ with std=2Å (see methods) and selected Neff=5 for the input MSA. Despite providing IF spin-label distance restraints, the predictions persisted in an OF state (**Fig. 5A**) which is the initial unconstrained prediction for PfMATE. While the RMSE to the target IF model decreased from 11.75Å to 9.07Å, AlphaFold2 failed to fold according to the input spin label constraints.

**Fig. 5:**
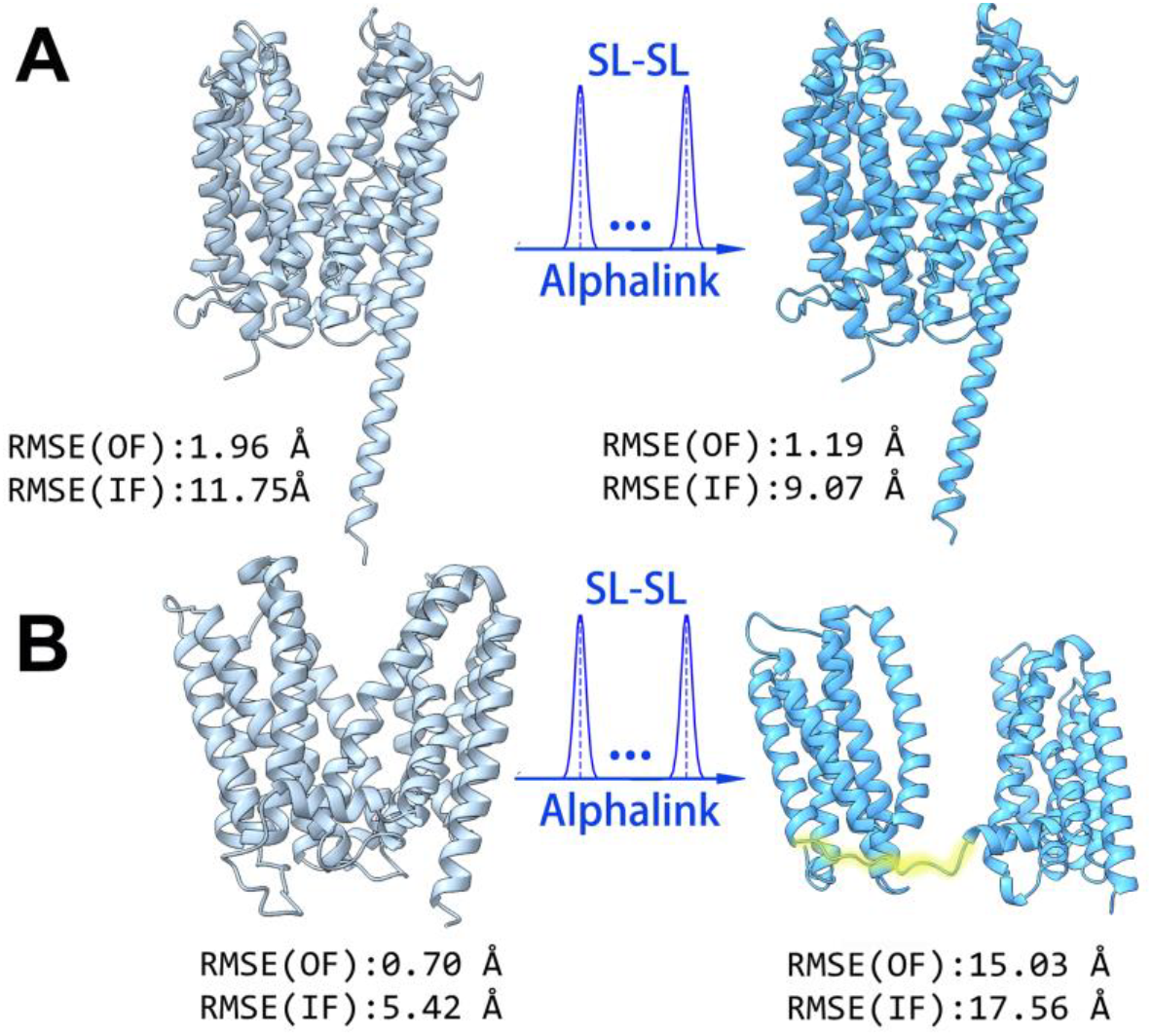
Alphalink prediction with spin-label distance distributions. **A**. PfMATE. **B**. LmrP(Highlighted region is from residue 192 to 197).

For LmrP (**Fig. 5B**), we generated input distances from spin label pairs reported in the paper^28^. When incorporating the IF spin label distances, the resulting model in **Fig. 5B** exhibited poor folding with an average predicted lDDT (pLDDT) score of only 64.35 in the central loop region (residue 192-197). Substantial deviations from both initial and target states were observed (with the RMSEs of 15.03Å and 17.56Å respectively). Structural inspection revealed the two helical subdomains were splayed far apart, suggesting the predictions were heavily biased by inconsistencies between the spin label and actual Cα-Cα distances.

### Refining AlphaFold2 with DEER Distance Distributions

The limitations uncovered above are not unexpected since AlphaLink^37^ was refined using alpha carbon distances and a distance range smaller than 42Å. The rotameric freedom of the spin label side chain introduces large uncertainties that probably exceed the conservative standard deviation range tested above. Therefore, we developed a new method, DEERFold, that fine-tunes the AlphaFold2 network using spin-label distance distributions. Capitalizing on the OpenFold^45^ platform, we incorporated DEER distance restraints into the AlphaFold2 network via the Evoformer module, supplementing the pre-existing co-evolutionary information (see **Fig. 1**). DEER data can be incorporated into the pair representation as the pairwise distance relationship between residue pairs, replacing template information. We then update the pair representation by adding the embedded DEER data to the original pair representation, resulting in a new pair representation that now includes the additional DEER distance information. The Evoformer block allows the MSA representation and pair representation to exchange information biased toward the DEER distogram data as described in **Fig. 1**. Notably, using only 6156 protein chains, which consist of approximately 4.7% of the full OpenFold training data, proved sufficient to fine-tune AlphaFold2 with DEER distance distributions and achieve reasonable accuracy, as shown below, for the purpose of this work.

To comprehensively evaluate DEERFold performance, we tested two different sets of distance inputs: experimental DEER data and simulated distance distributions. The experimental data, derived from previously published papers^27,28^, were processed in two ways: raw distributions determined from multi-Gaussian fitting of the DEER decays (**Experiment 1**) and a single-gaussian approximation (**Experiment 2**) as described in the Methods section. Simulated data were generated using the optimization method of Kazmier et al.^72^, with two sets created based on different helix length thresholds (**Simulation 1** and **Simulation 2**, see Methods section).

The results are shown in **Fig. 6** and **Fig. 7** for PfMATE and LmrP respectively. To ensure the robustness of our analysis, we generated 100 independent models for each target protein with different seed. We conducted tests using a shallow MSA (Neff = 5) and the full MSA for comparison. As noted previously^23^, MSA subsampling alone did not yield distinct conformational states for either transporters. For evaluation of the overall DEERFold prediction accuracy, we calculated the RMSD of Cα atoms between the predicted and target models.

### Experimental DEER distance distributions guide AlphaFold2 to the target conformation

The unconstrained PfMATE model at Neff = 5 closely matches the OF conformation (6GWH ^55^) (**Fig. 6A** model in light blue on the left), with a median RMSD of 0.87 Å, whereas the RMSD was 5.19 Å to the target IF state (6FHZ^55^). Including experimental spin label distance distributions in the prediction for both targets (denoted as Experiment 1 and Experiment 2, **Fig. 6C** for Neff = 5), highlight the robust performance of DEERFold. In Experiment 1, the predicted models showed a median RMSD of 2.11 Å to the target IF state, significantly lower than the 3.53 Å RMSD to the unconstrained OF conformation. Experiment 2, which employed approximated experimental data derived from Experiment 1, yielded a median RMSD to the target IF state of 1.32 Å, while the RMSD to the initial OF conformation increased to 4.71 Å, indicating a more substantial shift towards the target structure.

Increasing the MSA depth to full MSA tightens the range of predictions across all constraints sets (Fig. 6C). This tightening for the Experiment 1 data set yield predictions that exhibit a narrower IF conformation, with median RMSD values of 2.41 Å and 2.88 Å to the IF and OF states, respectively. In terms of TM score, the structural similarity remains high, with 93% of predictions achieving a TM-score > 0.9 to the IF state.

**Fig. 6:**
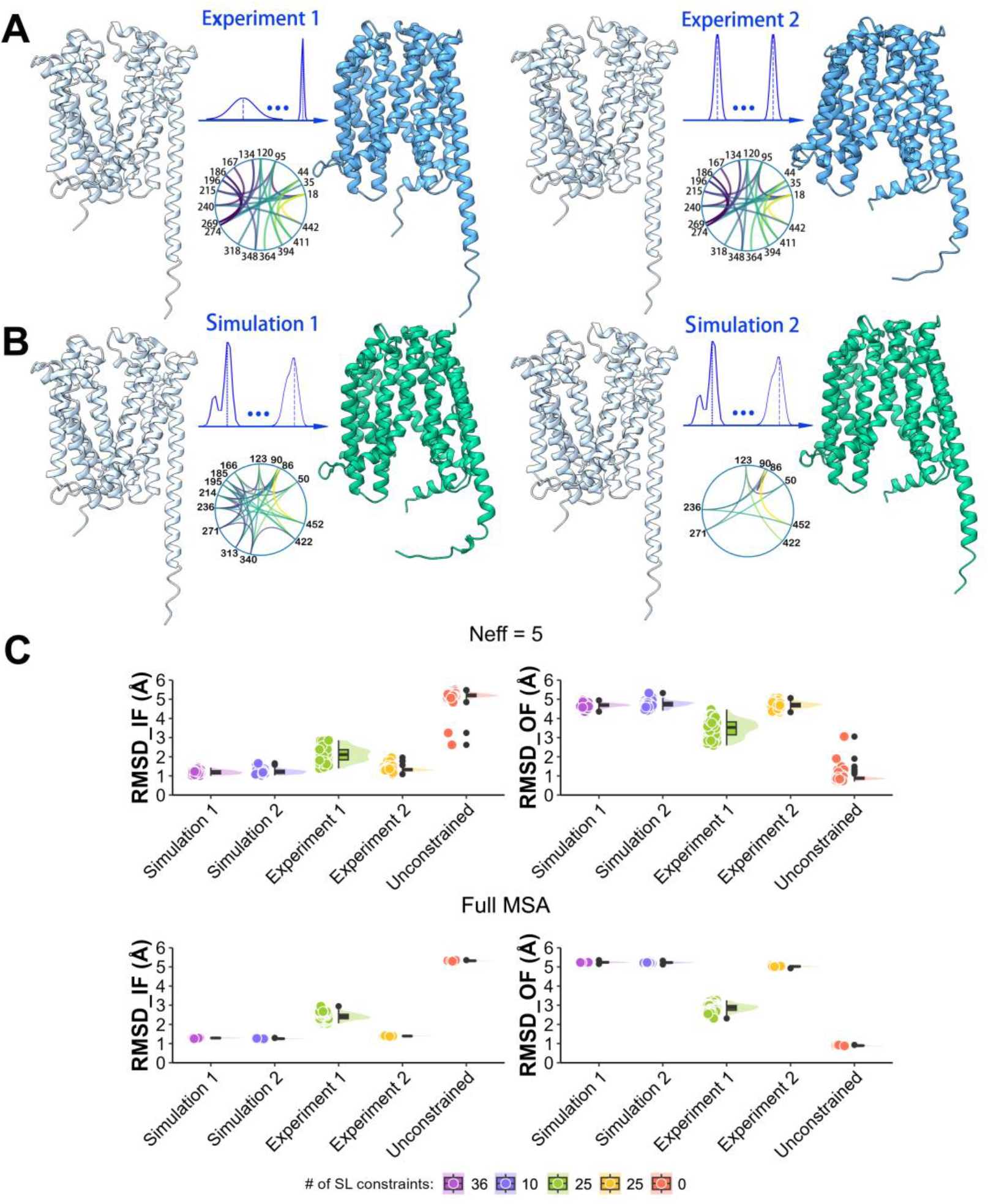
DEERFold predicted models of PfMATE with different sets of spin-label distances. **A**. Unconstrained DEERFold prediction which favors the OF conformation in light blue and the predicted DEERFold models from experimental constraint sets in deep blue. The DEERFold models were predicted using **Experiment 1:** raw experimental data (left); **Experiment 2:** approximated experimental data (right). **B**. Unconstrained DEERFold prediction in light blue and the predicted DEERFold models from two simulated constraints sets in deep green. **Simulation 1:** the larger set of simulated restraints with helix segments longer than 20 residues (left); **Simulation 2:** the reduced set of simulated restraints with helix segments longer than 30 residues (right). The detailed constraints are provided as chord diagram. **C**. RMSD between the predicted models for each constraint set and the referenced conformation states (IF/OF) at Neff = 5/ Full MSA.

For LmrP, the unconstrained prediction at Neff = 5 yielded an OF conformation. Using the raw experimental data (Experiment 1), yielded models with a median RMSD reduced from 5.56 Å (unconstrained set) to 1.52 Å to the IF state (AlphaFold2_prediction^57^). Conversely, the median RMSD to the initial OF state (6T1Z^56^) increased from 2.06 Å to 5.20 Å, as illustrated in **Fig. 7C**. Experiment 2 showed more pronounced switching, where the median RMSD to the IF state further decreased to 1.19 Å, while that to the OF state increased to 5.67 Å. Similar to PfMATE, a full MSA tightens the range of predictions across all constraint sets. However, both experimental sets for LmrP can “coerce” switching to the IF state, achieving RMSD below 1.5 Å. The overall trend suggests that deeper MSAs, while increasing prediction homogeneity, may in some cases override raw experimental distance constraints (Experiment 1), potentially limiting conformational diversity. However, this limitation can be overcome through idealized experimental restraints (Experiment 2), which maintain their ability to guide structural predictions even with deeper MSAs.

**Fig. 7:**
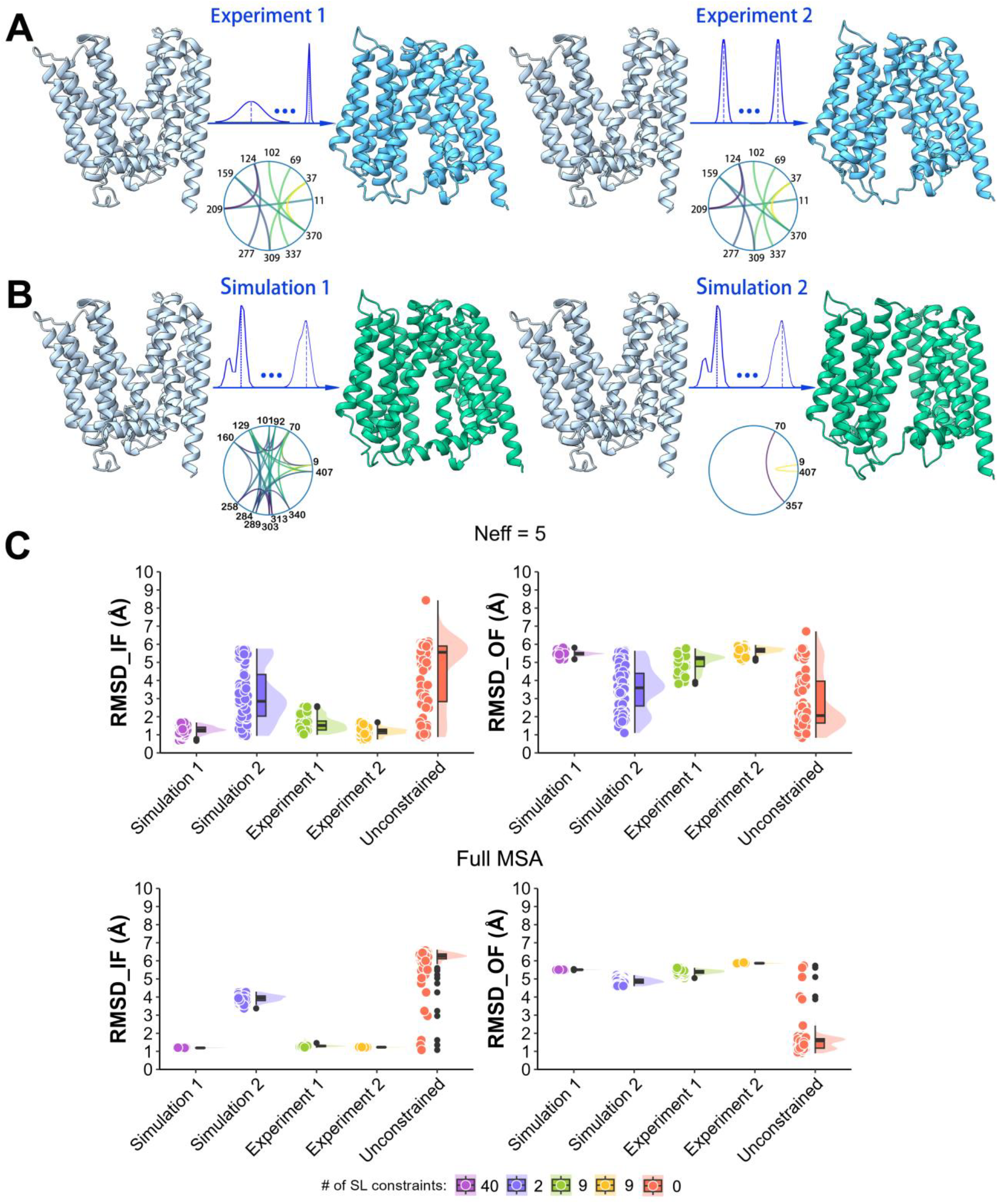
DEERFold predicted models of LmrP with different sets of spin-label distances. **A**. Unconstrained DEERFold prediction favors the OF conformation in light blue and the predicted DEERFold models from experimental constraint sets in deep blue. The DEERFold models were predicted using **Experiment 1:** raw experimental data (left); **Experiment 2:** approximated experimental data (right). **B**. Unconstrained DEERFold prediction in light blue and the predicted DEERFold models from two simulated constraints sets in deep green. The DEERFold model were predicted using **Simulation 1:** the larger set of simulated restraints with helix segments longer than 20 residues (left); **Simulation 2:** the further reduced set of simulated restraints with helix segments longer than 30 residues (right). The detailed constraints are provided as chord diagram. **C**. RMSD between the predicted models

### Optimized spin label pairs for conformational switching

As described in Methods, the simulated data were selected following the method of Kazmier et al.^72^ which optimize the information content by placing the spin label pairs at the ends of helix/beta strand segments. Since PfMATE and LmrP contain no beta strands, **Simulation 1** generated a larger set of simulated restraints with helix segments longer than 20 residues. As shown in **Fig. 6C** and **Fig. 7C** at Neff = 5, **Simulation 1** improved DEERFold’s performance for both transporters compared to the raw experimental set **Experiment 1**, with the lower median RMSD of 1.19 Å(PfMATE) and 1.28 Å(LmrP) to the target IF state.

We further tightened the criteria to include only helix segments with lengths ≥ 30 residues (**Simulation 2**), reducing the constraints to 10 for PfMATE and 2 for LmrP. Remarkably, even with this extremely sparse set of constraints, DEERFold successfully drove conformational switching for a significant subset of models from the initial OF state to the target IF state. As shown in **Fig. 6C** for PfMATE, all the DEERFold predicted models closely resembled the target state, with RMSD values of 1.22 Å between the predicted models and the target structures. Similarly, for LmrP (**Fig. 7C**), even with only 2 spin-label distance constraints, 58 out of 100 models approached the target state, with the RMSD values below 3 Å relative to the target structures. However, with 2 constraints at full MSA (**Fig. 7C**), DEERFold fails to fully switch to the target IF state from the initial unconstrained model, although the predictions show a trend toward the IF state, with a median RMSD value of 3.94 Å to the IF state compared to 4.86 Å to the OF state.

Comparing the performance across all distance sets, we observed that a limited number of spin label optimized distance sets were sufficient to guide DEERFold to the target conformation, with additional constraints showing no significant improvement in performance. Notably, DEERFold demonstrated robust performance with both simulated and experimental data. While the success with simulated constraints was expected, the comparable or even superior performance achieved with experimental data, particularly with approximated experimental constraints, highlights the practical utility of our approach. Furthermore, DEERFold showed remarkable efficiency in guiding models toward target conformations even with limited distance constraints, demonstrating its effectiveness in real-world applications where experimental constraints might be sparse.

### Application of DEERFold to the multidomain ABC transporter Pgp

Having established the performance of DEERFold on single domain membrane transporters, we tested its application to the more challenging multi-domain ABC transporter, Pgp or ABCB1. In addition to its helical transmembrane domain (TMD), Pgp has two cytoplasmic domains, the ATP binding cassettes or nucleotide binding domains (NBDs) that bind and hydrolyze ATP. The NBDs have a mixed helix/sheet architecture where these secondary structures tend to be few amino acids in length. Finally, Pgp was the subject of a DEER investigation into its ATP-dependent conformational dynamics yielding 24 distance distributions under different biochemical conditions ^74,75^. Since only two distinct conformations of Pgp have been determined to high resolution, we focused here on DEER data sets obtained under ligand-free (apo) conditions, and in the high energy post-hydrolysis intermediate trapped by Vandate following the hydrolysis of ATP (ADP-Vi conditions). The two conditions stabilize predominantly IF and OF conformations of Pgp respectively. For purpose of the comparison below, we focus on four structures of Pgp (7A65^59^, 7ZK7^58^,7ZK4^58^, and 6C0V^76^) representing two IF and two occluded/OF conformation as shown in Supplementary **Fig. 2**.

As shown in **Fig. 8A**, unconstrained DEERFold predicts narrow IF conformation of Pgp, with a median RMSD value of 3.41 Å compared to experimental structure (7A65^59 54^), and a median RMSD value of 6.29 Å and 6.24 Å compared to the OF structures (7ZK4^54^, 6C0V^76^). Including the raw distance distributions (Experiment 1), corresponding to the ADP-Vi intermediate, reduces the median RMSD to the OF structures to 3.25 Å(7ZK4) and 3.30 Å(6C0V). Similarly, Experiment 2 yielded comparable results with median RMSD values of 3.25 Å and 3.29 Å for 7ZK4 and 6C0V, respectively (see **Fig. 8A** and **Fig. 8C**). In contrast, predictions with the Apo dataset yielded conformations with a wider intracellular opening than both reference IF structures (7A65 and 7ZK7). Models predicted with Experiment 1 data set have a median RMSD to the IF-wide structure (7ZK7) of 4.90 Å down from 6.30 Å while simultaneously increasing the RMSD from 3.41 Å to 4.90 Å for the IF-narrow structure (7A65). Experiment 2 induced an even more pronounced opening, with the median RMSD of 5.34 Å to the IF-wide structure(7ZK7), and the median RMSD to the IF-narrow structure further increasing to 11.01 Å (see **Fig. 8B** and **Fig. 8C**).

**Fig. 8:**
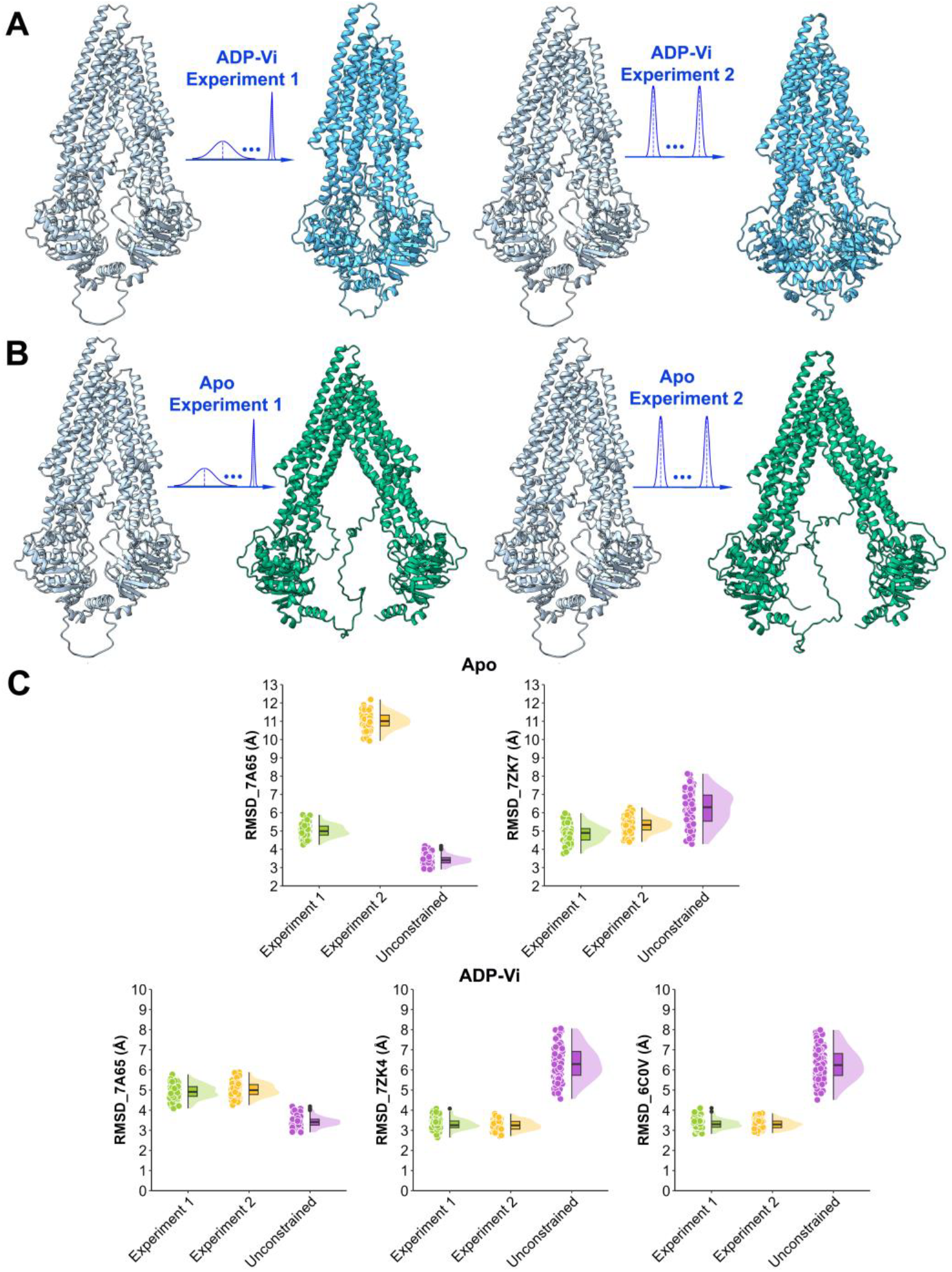
DEERFold predicted models on the Pgp with different sets of spin-label distances. **A**. Unconstrained DEERFold prediction favors the IF-narrow conformation in light blue and the predicted DEERFold models from two ADP-Vi constraint sets are shown in deep blue. The DEERFold models were predicted using **Experiment 1:** raw experimental data (left); **Experiment 2:** approximated experimental data (right). **B**. DEERFold models from two Apo constraint sets are shown in deep green. The DEERFold models were predicted using **Experiment 1:** raw experimental data (left); **Experiment 2:** approximated experimental data (right). **C**. RMSD between the predicted models for ADP-Vi/Apo and the referenced conformation states (IF/OF) at Neff = 10.

### Sparse distance constraints can fold Pgp into the target conformation

Finally, we evaluated whether DEERFold can achieve conformational switching with simulated DEER constraint sets that are smaller than the reported experimental throughput^74,75^. We selected a simulated distance set (Simulation 2) from the target state(7ZK7), comprising 54 pairs of restraints derived from helix segments ≥ 30 residues and beta strand segments ≥ 5 residues. From this set, we randomly selected 8 constraints as inputs for DEERFold and evaluated the models using TM-scores against both reference structures (7A65 and 7ZK7). The comparison in terms of TM-score, shown in **Fig. 9B**, indicates a trend of the DEERFold prediction transitioning from the initial IF-narrow-like conformation (similar to7A65). Notably, 293 out of 500 models successfully switched to the target wide IF state (7ZK7), with a TMscore ≥ 0.9 relative to 7ZK7(blue dots in **Fig. 9B** right plot). The notable success rate (58.6%) with randomly selected constraints indicates that conformational switching can be achieved at a subset of possible spin label pairs, instead of exhaustive measurements. For comparison, **Fig. 9A** and **Fig. 9B** (left plot) presents the conformational sampling of Pgp under unconstrained conditions and full set of constraints from the target wide IF state (7ZK7). With the full constraint set (54 pairs), all predictions successfully transitioned to the wide state (Fig. 9B left plot). In contrast, unconstrained predictions predominantly favored the narrow IF state, with a mean TM-score of 0.93 to 7A65. However, the unconstrained predictions showed some structural diversity, as highlighted by two distinct conformations (yellow dots in Fig. 9C) that exhibited the largest deviations among all models. Two representative models showing chain thickness based on RMSF values relative to the narrow IF state (7A65) are presented in Supplementary **Fig. 3**.

**Fig. 9:**
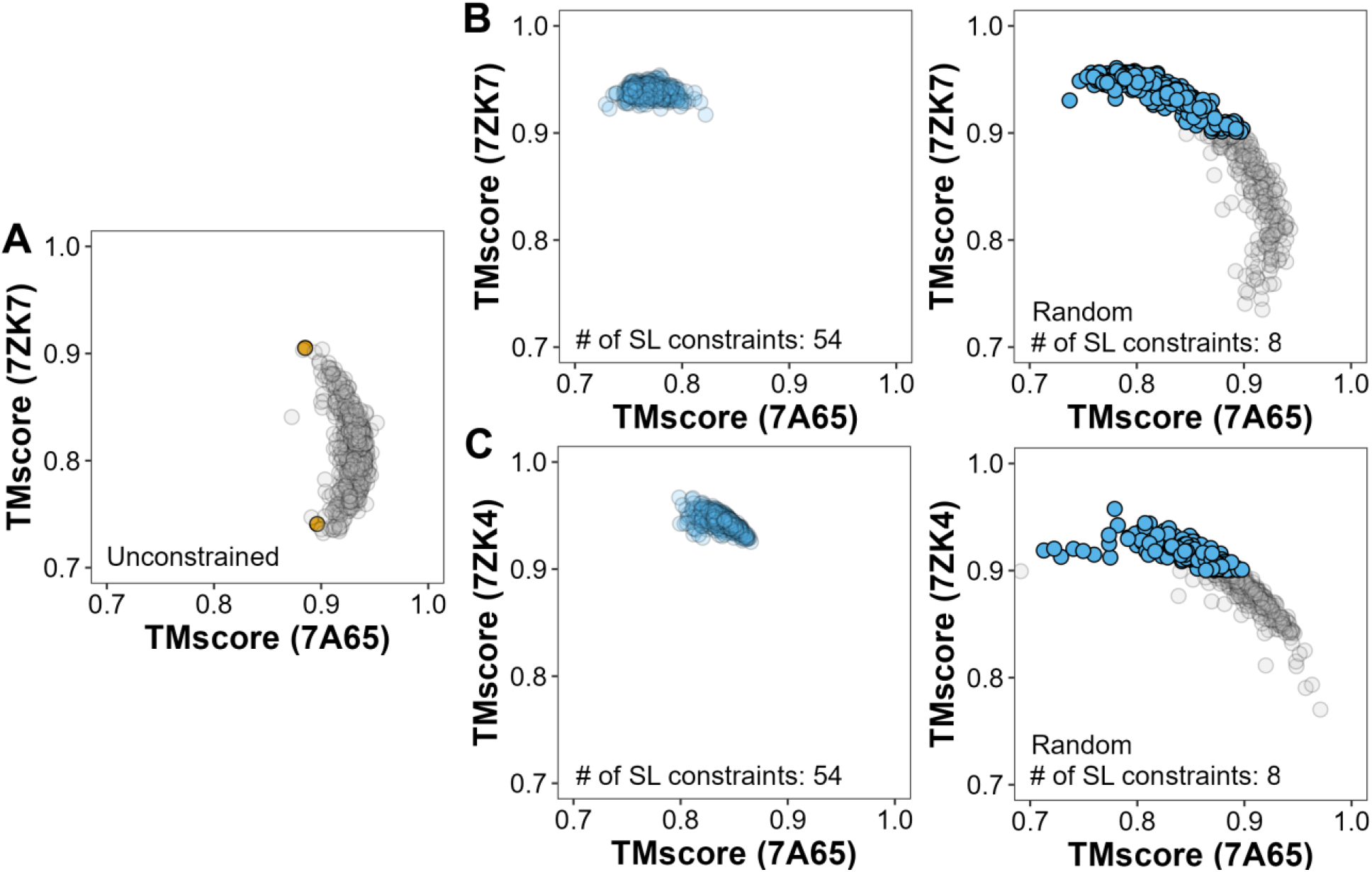
Conformational sampling of Pgp using DEERFold. **A**. Unconstrained predictions of Pgp (yellow dots with two distinct conformations among all unconstrained predictions). **B**. Predictions constrained by the simulated distance set (Simulation 2) derived from the target state(7ZK7). **Left**: using the full set constraints of 54 pairs. **Right**: 8 randomly selected simulated DEER distance distributions from the full set (blue dots with TM score≥ 0.9 to 7ZK7). **C**. Predictions constrained by the simulated distance set (Simulation 2) derived from the occluded/OF state(7ZK4). **Left**: using the full set constraints of 54 pairs. **Right**: 8 randomly selected simulated DEER distance distributions from the full set (blue dots with TMscore≥ 0.9 to 7ZK4). All panels show the TM score values of predicted conformations relative to known narrow IF (7A65) and wide-IF (7ZK7) / occluded-OF (7ZK4) states.

Given that unconstrained predictions favor the narrow IF state, we further tested DEERFold with simulated OF state (7ZK4) constraints. All models successfully transitioned to the OF state with the full constraint set, while 187 out of 500 predictions showed partial folding toward the OF state under randomly selected constraints.

### Earth mover distance score for identification of alternative conformations

In the absence of experimental structures, it will be challenging to evaluate the DEERFold results to determine whether the distance constraints are sufficient for conformational switching. To overcome this obstacle, we evaluated an alternative method that relies on the EMD between the distance distributions of the predicted model compared to the input distance distributions. To illustrate this method, we generated a set of constraints for the target conformation of LmrP, PfMATE, Pgp, following the method of Kazmier et al.^72^ (referred to as Simulation 2 set in Methods section). We chose shallow MSA(Neff = 5 for PfMATE and LmrP; Neff = 10 for Pgp), and randomly selected subsets of simulated constraints (2 out of 40 for LmrP; 25 out of 36 for PfMATE; 8 out of 54 for Pgp) from the target conformation. These sets were then used in DEERFold to generate 500 models for each target.

To analyze the generated models, we performed Principal Component Analysis (PCA) with ProDy^77^. Variation along PC1 was plotted against the EMD values followed by K-means clustering (via the sklearn library^78^) along PC1. The centroids (cluster-representative models selected by structure alignment tool Foldseek^79^) are marked as red dots in **Fig. 10**. For each of the targets, we observed that different clusters generally represent distinct conformations, each with a different EMD value. As shown in **Fig. 10**A for LmrP, the PC1 values clustered into three clusters, demonstrating a conformational switch from the OF to the IF structure as the EMD distance decreases. ^53^The yellow cluster centroid shows high structural similarity to the reference OF structure (6T1Z), with an RMSD of 0.81 Å and a TM score of 0.99. The intermediate cluster centroid (shown in red) displays partial characteristics of both states, with metrics relative to the OF state (RMSD = 3.91 Å, TM-score = 0.80) and the IF structure (AlphaFold2_prediction53; RMSD = 2.69 Å, TM-score = 0.89). The blue cluster centroid, which has the lowest EMD distance, aligns closely with the IF structure (RMSD = 1.73 Å, TM-score = 0.96). Similarly, for PfMATE, illustrated in **Fig. 10B**, the PC1 values are roughly clustered into three parts. The first centroid in yellow has a RMSD of 1.16 Å to the reference OF structure (6GWH), and a TM score of 0.98. The metrics of the gray cluster centroid (shown in red) suggest an intermediate more similar to the OF (RMSD = 2.41 Å, TM-score = 0.92) than the IF structure (6FHZ; RMSD = 2.87 Å, TM-score = 0.88). Remarkably, the blue cluster centroid, which has the lowest EMD distance, aligns more closely with the IF structure (RMSD = 2.03 Å, TM-score = 0.93).

**Fig. 10:**
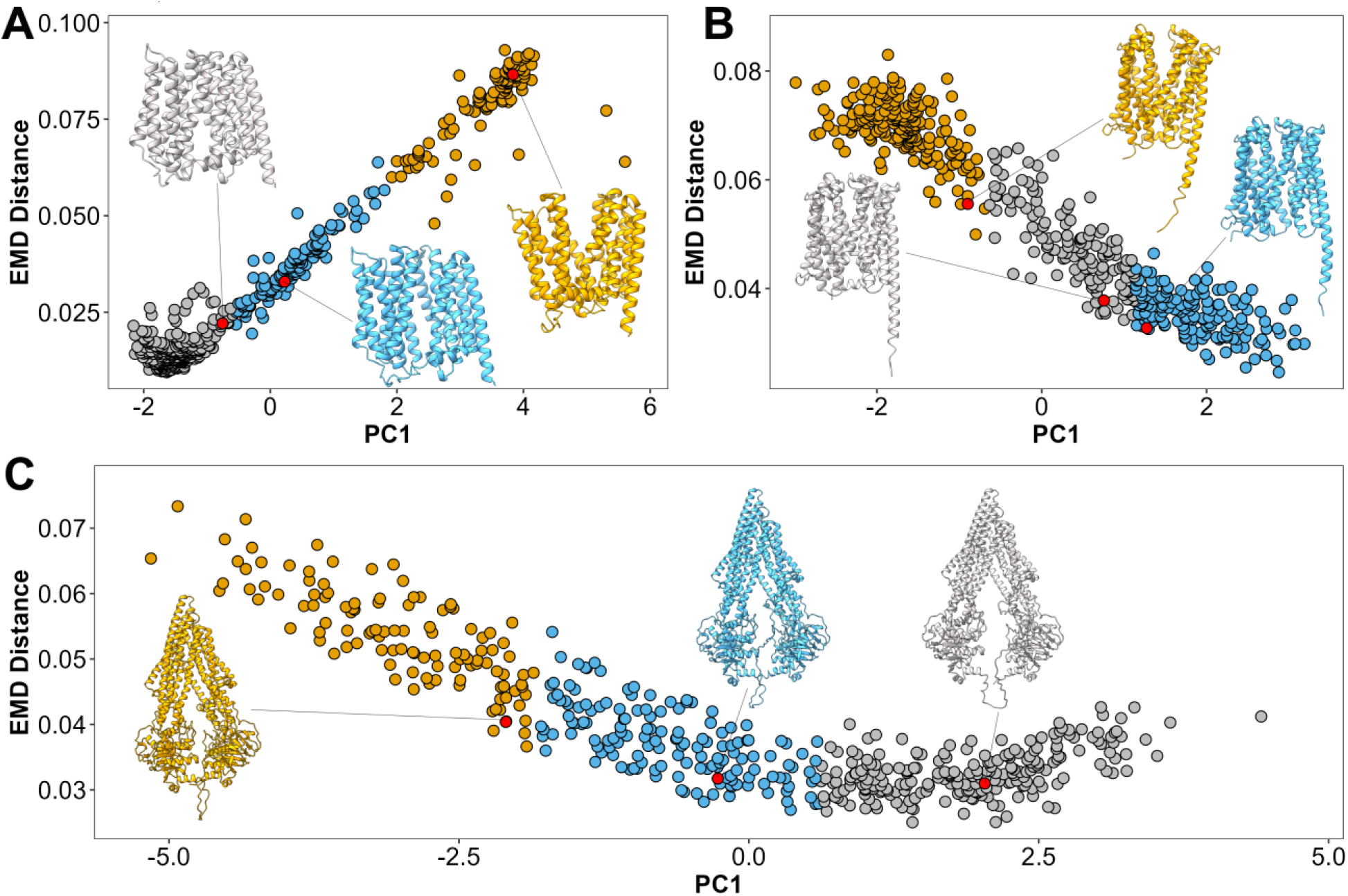
Evaluation of DEERFold predictions with PCA and EMD distances. **A**. LmrP: PC1 values clustered into three. Red dots indicate three cluster centroids. **B**. PfMATE: PC1 values form three main clusters. Red dots indicate three cluster centroids. **C**. Pgp: PC1 values divide into three categories based on EMD values, showing gradual opening of IF conformations. Red dots indicate three cluster centroids.

The results for Pgp were particularly interesting (**Fig. 10C**). We observed that the structural diversity represented by the clusters along PC1 represents IF conformations with gradual opening with RMSDs decreasing from 5.38 Å to 2.93 Å and increasing TMscores from 0.84 to 0.95 relative to the reference IF structure(7ZK7). All results demonstrate a strong correlation between EMD scores and PC1 values, indicating the EMD score as an effective metrics for model selection. This correlation is further bolstered by the relationship between EMD scores and TM-scores shown in Supplementary **Fig. 4**. Lower EMD distances, indicating better agreement with input distance constraints, consistently correspond to higher TM-scores relative to the target structure.

### General performance of DEERFold

To demonstrate the generality of the method and extend it to globular proteins, we tested DEERfold on other protein targes including Adenylate Kinase (AK), Ribose Binding Protein (RBP), and a set of membrane protein targets, PfMATE, LmrP and Pgp, MCT1, STP10, LAT1, ASCT2, E. coli SemiSWEET and DgoT. The structure of each of these targets was determined in at least two conformations (details in Methods section). The unconstrained prediction by DEERFold was set as the initial conformation making the other experimental structure the target conformation. We used DEER simulated data by the Kazmier method^72^ for evaluation. For better visualization, we set Neff = 5 to generate 15 DEERFold predictions for each. Complete results are shown in **Fig. 11**.

DEERFold drives the models toward the target states across a diverse set of protein targets with limited number of distance constraints (≤ 25). As shown in Fig. 11A, the constrained predictions in blue successfully capture significant conformational transitions from their unconstrained starting states in brown. The quantitative analysis in Fig. 11B reveals that DEERFold predictions (red dots) consistently achieve higher TM scores with respect to the target structures compared to unconstrained predictions (purple dots). Notably, even with minimal spin label constraints (e.g., DgoT and E. coli SemiSWEET with only 2 constraints), DEERFold could successfully guides part of the predictions toward the target conformations. This remarkable success established the generality of DEERFold.

**Fig. 11:**
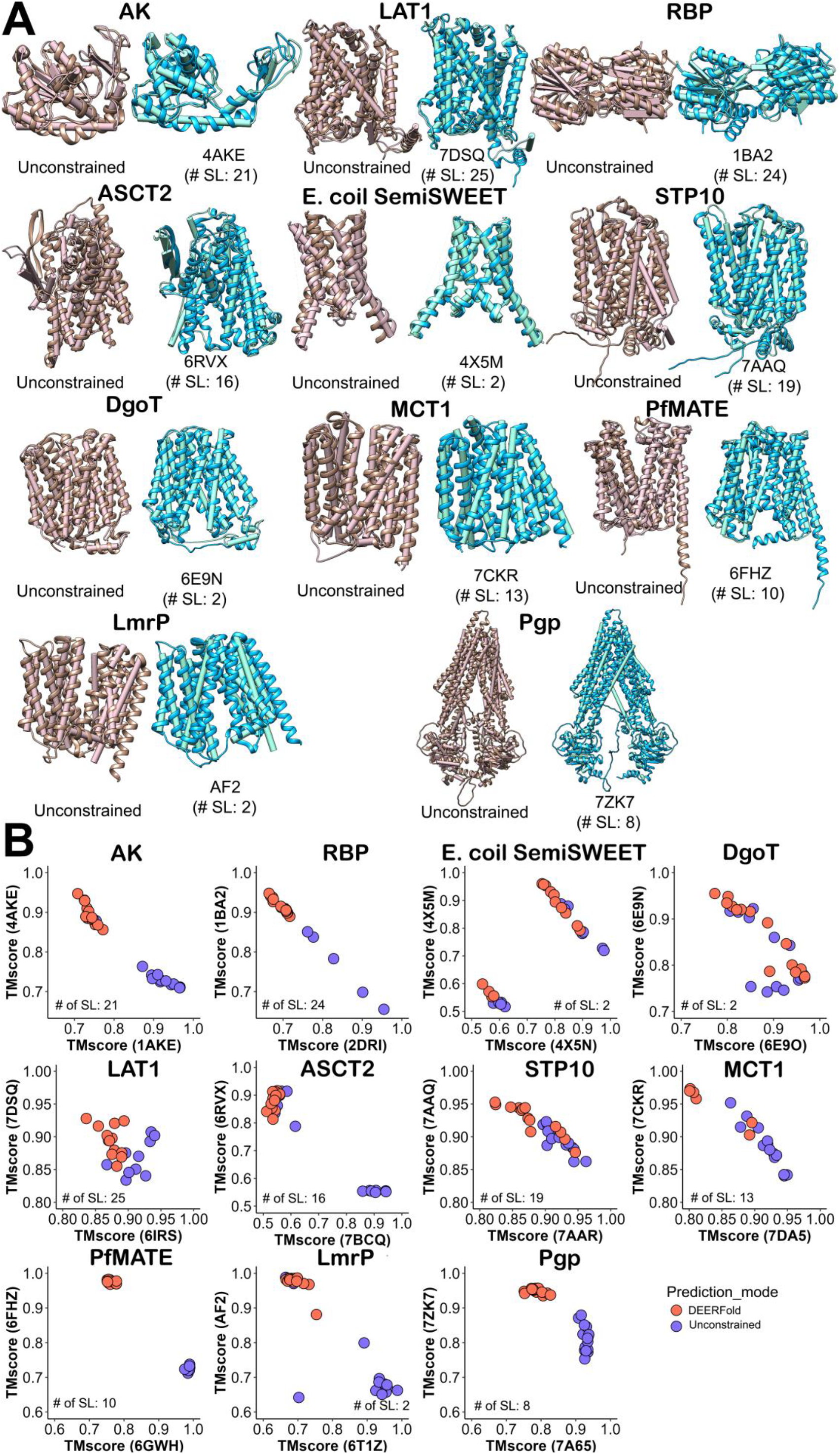
Performance of DEERFold on models trained by spin labels derived from beta strand (DEERFold_strand) and helix (DEERFold_helix). **A**. Best prediction results for DEERFold, superimposed with the experimental structure. Blue:Distance constrained DEERFold prediction, Brown: Unconstrained Prediction. DEER distance constraints are derived from the target state of each protein. **B**. Evaluation results on protein targets presented in terms of TMscore.

## Discussion

Multiple methods have been advanced to fold proteins or predict their structures with experimental constraints. While the most successful methods have been in the field of NMR spectroscopy^80^, the problem becomes more challenging if the restraints are sparse. Another layer of complexity is introduced when the constraints are derived from probes that are attached to the backbone via flexible side chains. Previous attempts to model sparse spin label constraints using physics-based forcefields^41,81^ or knowledge-based methods^82,83^ such as Rosetta had limited success.

Here, we developed a general strategy to integrate distance distributions into the prediction of AlphaFold2 network architecture. We showcase its success with a novel approach called DEERFold that harness the spatial constraints encoded by spin label distance distributions to generate conformational distributions. The hypothesis driving this work is that the more robust implicit energy function of AlphaFold2, as manifested by the accuracy of its models, will overcome the sparseness and the uncertainties of the probe-based constraints. Although current methods to simulate spin label distance distributions^41–44^ overstate their widths, AlphaFold2 was able to overcome this limitation as manifested by the precision of our results. during training.

In addition to establishing the conceptual feasibility of supplementing AlphaFold2 prediction with probe-based distance distributions, the work presented here investigates the parameters that affect its performance including the depth of the MSA and number and width of distance distributions. Remarkably, an approximate representation of the experimental data, where the width is set at 2 Å, performed equally well with the unmodified distributions. This finding may have direct significance to the experimental throughput since obtained an accurate width is typically a bottleneck in the DEER experiment. Overall, the performance of DEERFold reflects the robustness of the underlying AlphaFold2 network enabling relatively sparse, uncertain constraints to drive the prediction of alternative conformations. From an intuitive perspective, the constraints appear to redirect AlphaFold2 to alternative hypotheses which are then refined into models. We illustrated the generality of DEERFold on multiple classes of proteins.

Two critical parameters shape the DEERFold inference. As expected, larger MSA or Neff can bias DEERFold to the default prediction reducing the impact of the distance constraints. The Neff of MSAs was set to less than 25 to reduce the bias without compromising the quality of the models. Neff value during inference is likely to depend on the target and should be varied. The information content of the distance constraints emerges as a critical parameter affecting the performance of DEERFold. Although the experimental constraints were able to drive DEERFold to alternative conformations, some of the spin label pairs were selected in the absence of high-resolution structures. Hence, a pair selection method that maximizes the information content of the constraints, previously introduced in the context of ROSETTA, increased the quality of the predicted models. It is possible that a neural network approach can further optimize the selection of spin label pairs.

Approximating the conformational space by PC1 in a PCA, we showed that low EMD between the input and predicted distance distributions can identify clusters of models of the target conformation. Although within each cluster, there are structural differences, as evidenced by similar EMD values but varying PC1 values, the general trend indicates that clusters with lower EMD distances are more similar to the target conformational state. Further analysis, such as putty analysis on the top 5 models with the lowest EMD values within each cluster, reveals that structural differences typically occur in unstructured regions, as shown in Supplementary **Fig. 3**. The successful application of DEERFold suggests that it can be extended to incorporate other types of experimental data, such as nuclear magnetic resonance (NMR), fluorescence resonance energy transfer (FRET), hydrogen-deuterium exchange (HDX), and cross-link mass spectrometry (XL-MS). DEERFold provides a general training platform for these complementary techniques to integrate into the AlphaFold2 system, enabling the incorporation of more valuable information about protein structure, dynamics, and interactions, which can significantly enhance the accuracy, reliability, and applicability of protein structure prediction.

## Supporting information

Supplementary doc

## Contribution

TW and HSM designed the research. TW carried out the research. TYK curated the data for training and inference. BB and RAS provided critical feedback during the design of the research and edited the manuscript. TW and HSM wrote and edited the manuscript.

## Notes

### Competing Interest Statement

The authors have declared no competing interest.

