## Supplementary doc for "Modeling Protein Conformations by Guiding AlphaFold2 with Distance Distributions. Application to Double Electron Electron Resonance (DEER) Spectroscopy"

Tianqi Wu[1], Richard A. Stein[1,2], Te-Yu Kao[2,3], Benjamin Brown[1,4], and Hassane S. Mchaourab[1,2*]
[1]Center for Applied AI for Protein Dynamics, [2]Department of Molecular Physiology and Biophysics, [3] Department of Chemistry, and [4]Department of Pharmacology, Vanderbilt University, Nashville, TN, USA

*

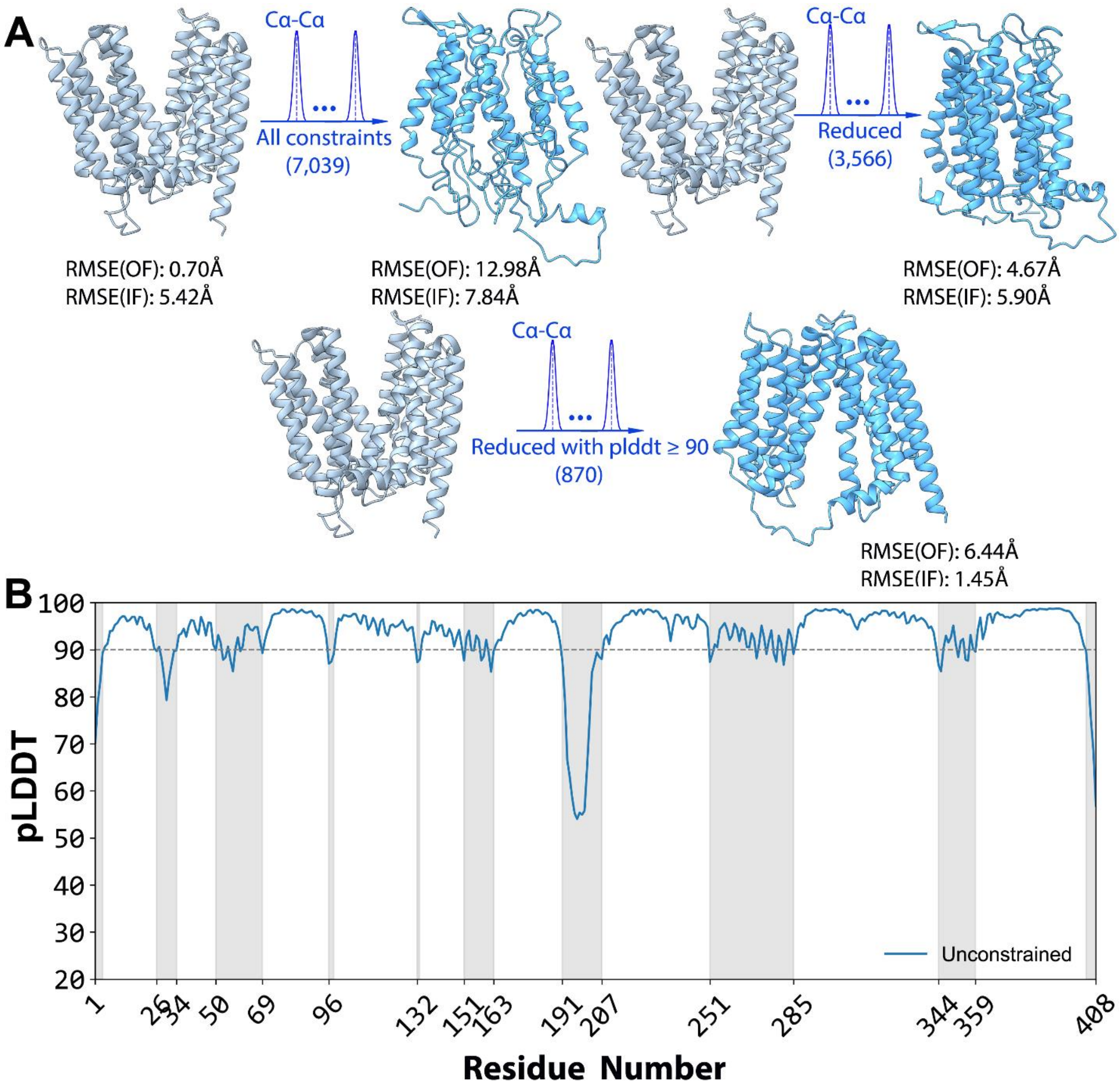


**Supplementary Fig. 1**: **Impact of distance constraints on AlphaLink prediction of the target conformation of LmrP**. **A.** The initial unconstrained AlphaFold2 prediction (light blue) favors the outward-facing (OF) conformation. Subsequent predictions using different constraint sets are shown in deep blue: **Set 1** employs all Cα-Cα distances(top left), yielding a model with RMSEs of 12.98Å to the default OF structure (AlphaFold2_prediction) and 7.84Å to the target IF structure (6T1Z); **Set 2** utilizes the subset of 3,566 distances from regions with structural differences (top right); **Set 3** implements an optimized subset of 870 distance constraints derived from high-confidence regions (pLDDT ≥ 90). The **Set 3** constraints successfully guided AlphaFold2 towards the target IF conformation. **B.** Per-residue pLDDT (predicted Local Distance Difference Test) score distribution for the unconstrained model. Gray regions indicate low-confidence areas (pLDDT < 90), corresponding to residue ranges [1,4], [26,34], [50,69], [96,98], [132,133], [151,163], [191,207], [251,285], [344,359], and [404,408], which were filtered out for the optimized constraint set in **Set 3**.

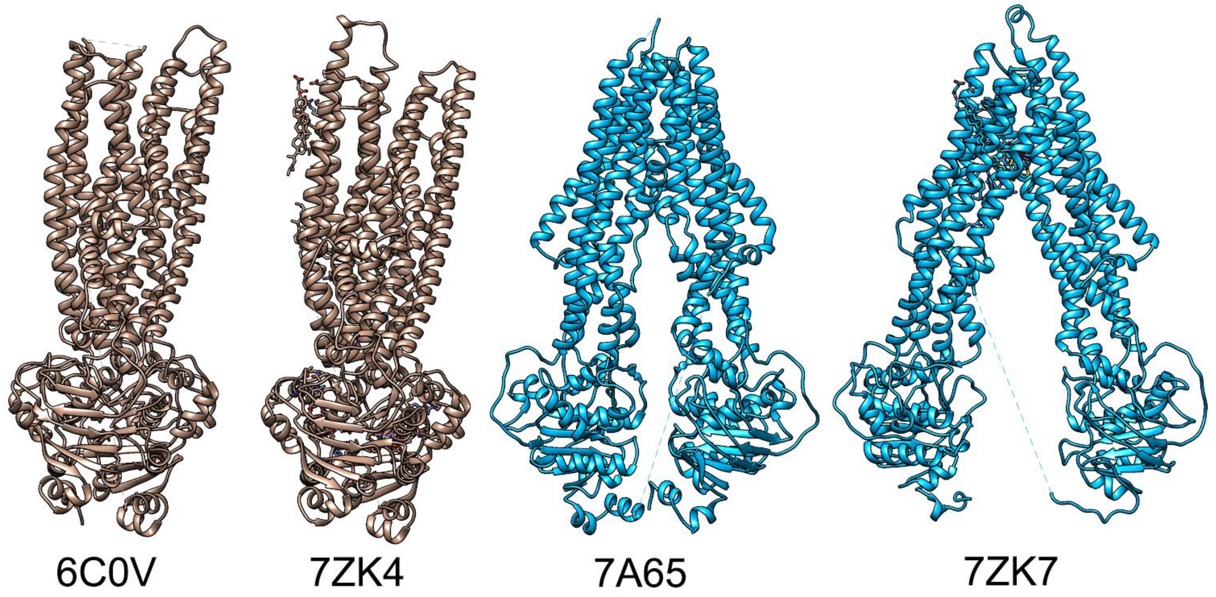


**Supplementary Fig. 2**: Comparison of native structures representing four conformational states of P-glycoprotein (Pgp). Structures shown are derived from PDB entries 60CV(Occluded-OF), 7ZK4 (Occluded-OF), 7A65 (IF-narrow), and 7ZK7 (IF-wide).

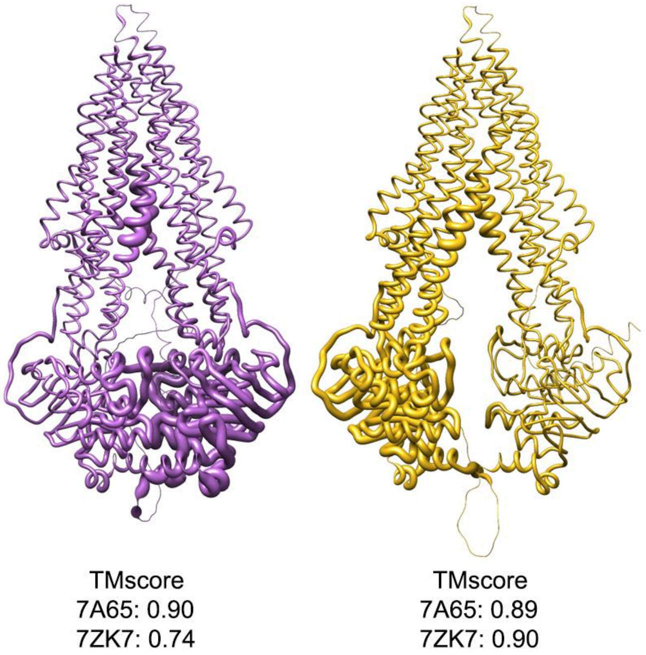


**Supplementary Fig. 3**: Structural differences in unconstrained DEERFold predictions for Pgp.Two representative models showing chain thickness based on RMSF values relative to the narrow IF state (7A65).

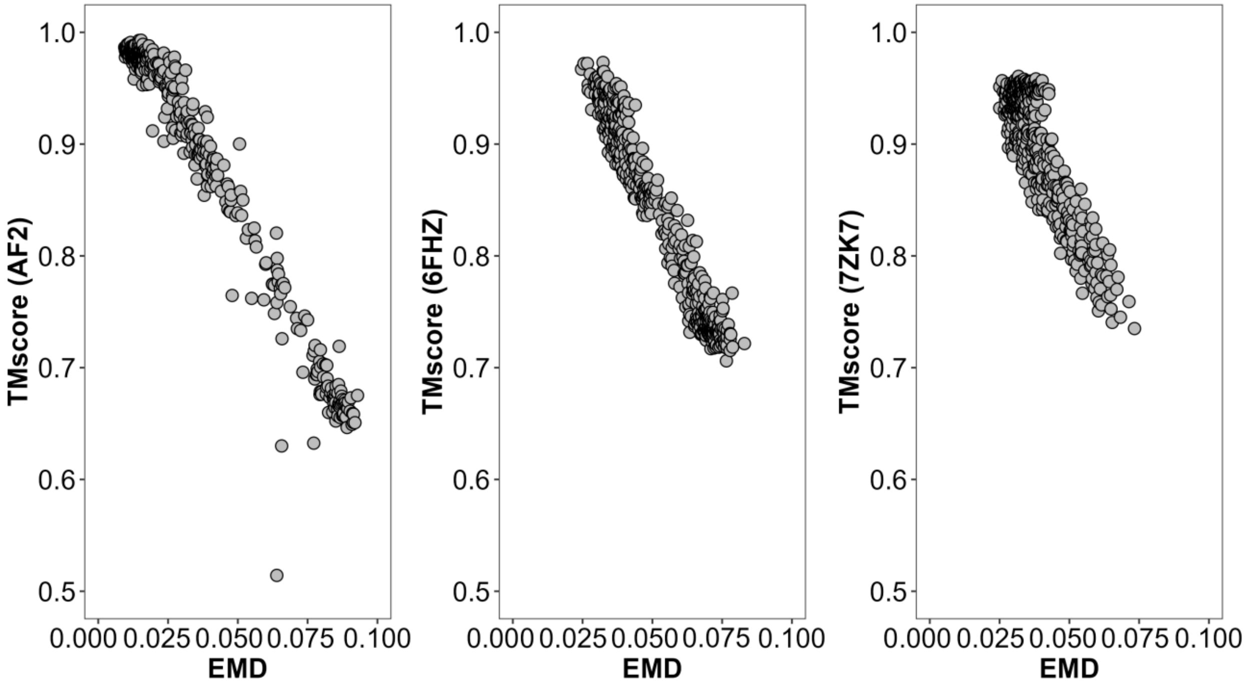


**Supplementary Fig. 4**: EMD scores versus TM-scores for 500 predicted models for LmrP(left), PfMATE(middle), and Pgp(right).
